# Benchmarking AlphaFold3-like Methods for Protein-Peptide Complex Prediction

**DOI:** 10.1101/2025.03.09.642277

**Authors:** Fan Zhou, Shangwei Guo, Xiangda Peng, Shuhan Zhang, Cuicui Men, Xiaobo Duan, Guoliang Zhu, Zechen Wang, Weifeng Li, Yuguang Mu, Liangzhen Zheng, Haibin Liu, Sheng Wang

**Affiliations:** Shanghai Zelixir Biotech Ltd, Shanghai, China; Dong’e Ejiao Co.,Ltd., Shandong Provincial Key Laboratory of Gelatine Medicines Research and Development, Liaocheng, Shangdong, China; School of Physics, Shandong University, Jinan, Shandong, China; School of Biological Sciences, Nanyang Technological University, Singapore; Shenzhen Zelixir Biotech Ltd, Shenzhen, Guangdong, China

**Author notes:** Corresponding Authors: Haibin Liu,; Sheng Wang. The authors contribute to the research equally (Fan Zhou, Shangwei Guo).

## Abstract

Protein-short peptide interactions are central mechanisms regulating signal transduction, chaperone functions, and drug targeting design. However, the precise prediction of their complex structures has long been constrained by the limited accuracy of traditional methods. The breakthrough of AlphaFold2 resolved the challenge of predicting protein monomer structures. Its derivative method, AlphaFold2-multimer, surpassed traditional docking approaches in predicting protein complexes, yet its accuracy remains suboptimal. AlphaFold3 and its replication models, such as Protenix, Chai-1, and Boltz-1, have further improved prediction accuracy for protein-protein interactions. To evaluate the performance of these methods in protein-short peptide structure prediction, this study conducted predictions and comprehensive comparisons on two benchmark datasets of protein-short peptide complexes. One dataset did not intentionally exclude structures from the training set of these methods, while the other did. For the former, the results demonstrate that the new-generation structural method significantly increased the success rate under stringent criteria (Fnat≥ 0.8) to 70-80%, compared to AlphaFold2-multimer (53%), with Protenix achieving the highest accuracy at 80.8%. For the second dataset, AlphaFold2-multimer showed consistent performance, but the success rates of the new-generation methods all dropped significantly (to 40-56%). Our analysis suggests that the main challenge for the performance decline comes from the accurate prediction of binding sites. For all datasets, A multi-method combination strategy enabled a higher-quality prediction success rate compared to the individual models, highlighting synergistic advantages among methods. Notably, inter-model consistency analysis provided an efficient metric for selecting optimal predictions.

## Introduction

Protein-short peptide interactions are critical to numerous biological processes. For instance, these interactions can directly participate in the activation or inhibition of signal transduction pathways and regulate key proteins within signaling pathways, thereby modulating the overall activity of the pathway [1]. Some short peptides act as molecular chaperones, assisting in the proper folding of proteins during synthesis [2]. For example, specific short peptide regions of heat shock proteins can recognize and bind to non-native structural regions of other proteins, preventing misfolding or aggregation and ensuring the formation of biologically active three-dimensional structures [3]. Furthermore, short peptide drugs enable specific binding to target proteins, enhancing therapeutic efficacy and selectivity while minimizing side effects [4, 5]. Accurate structural modeling of protein-short peptide complexes serves as a fundamental basis for studying these interactions. To this end, a series of methods have been developed, including traditional approaches employing empirical scoring functions (e.g., AutoDock-CrankPep [6], HADDOCK [7], RosettaDock [8]) and deep learning-based methods. [9–13].

AlphaFold2 [14], a deep learning-based method developed by DeepMind, enables accurate prediction of protein monomer structures. Subsequently, this approach was extended to complex prediction, demonstrating superior accuracy in molecular docking compared to traditional methods [15, 16]. AlphaFold3 (AF3), the next-generation iteration of AlphaFold2, incorporates multiple improvements [17]. Its Pairformer module reduces reliance on multiple sequence alignments (MSA) and emphasizes information from the target protein sequence itself. Additionally, AF3 introduces a diffusion model to replace the original iterative structural optimization, generating structures through a forward noise-adding and reverse denoising process. These advancements enhance AF3’s accuracy in predicting protein and complex structures while enabling the modeling of more diverse biomolecular interactions.

Chai-1, a replication of AF3, retains the core architecture and methodology of AF3. It has demonstrated performance comparable to AF3 in multiple benchmarks [18]. Boltz-1 represents the first fully commercialized open-source model. While inspired by AF3, it introduces innovations in model architecture, data curation, training, and inference workflows. For example, novel algorithms enable more efficient and robust MSA pairing, structural cropping during training, and conditional predictions based on user-defined binding pockets [19]. Protenix, another replication and refinement of AF3, is fully open source, providing comprehensive training code and datasets to facilitate broader accessibility and customization for researchers [20]. HelixFold3 further incorporates insights from the AF3 publication to replicate its advanced functionalities [21]. Collectively, we categorize these methods as **new-generation methods**.

The new-generation modeling approaches exhibit significant improvements over AlphaFold2-multimer (AF2m) in predicting protein-protein interactions. However, their specific advancements in protein-short peptide modeling remain under-explored. To address this gap, we evaluated AF3, Chai-1 (w/o MSAs), Boltz-1, and Protenix using two benchmark test sets in this work. The first test set comes from reference [22], including 99 protein-peptide complexes from PDB. The publication time of these complex structures was not used as a screening indicator to simulate the real application scenario - we do not need to care about whether the sequence is in the training set when applying these methods. The second test set includes 91 new protein-peptide complexes from PDB, which have not been included in the training set of any method involved in this work, reflecting the ability to predict new structures of these methods.

Our results reveal that the success rate of these methods under stringent criteria (Fnat≥ 0.8) reaches 70–80%, markedly surpassing AF2m’s 53%. Notably, combining multiple models further elevates the success rate to 90%. However, for the second test set, under the same criteria, the success rate of the new generation methods dropped significantly to 40%-60%, while that of AF2m remained unchanged at 55%. This suggests to some extent that the new generation methods have the risk of overfitting in protein-peptide modeling. We also explored the ability of various indicators, such as pLDDT, ipTM, RMSD, the number of interactions between peptides and protein receptors (#PPI) and model consistency in selecting the best prediction from multiple methods or models. Regardless of the first or the second test set, these indicators, especially model consistency, are helpful to identify which is the better predicted conformation.

## Results and Discussion

We use the Fraction of native contacts (Fnat)[23] to evaluate the quality of the model. Fnat is an important index to quantify the prediction accuracy of protein - protein docking models. It is defined as the ratio of the number of native interface contact residue pairs retained in the predicted model to the total number of native complex interface contact residues in the experimental crystal structure. Specifically, native interface contact pairs are determined by the set of atom pairs with heavy atom distances ≤ 5A in the crystal structure of two proteins (or protein - ligand). The value range of this index is [0,1]. The closer the value is to 1, the higher the compatibility of the model with the native structure’s interface. In the CAPRI (Critical Assessment of Predicted Interactions) evaluation system, Fnat ≥ 0.5 is usually considered to have medium accuracy, and Fnat ≥ 0.3 is usually considered to have acceptable accuracy [24, 25]. We use the criteria Fnat≥0.8 as the high accuracy in this work. Unless otherwise mentioned, the prediction success mentioned below means Fnat≥0.8.

According to stringent criteria(Fnat≥0.8), we found that all methods demonstrated good performance accuracy for the first test set, with improvements from 53% for AF2m to 70-80%. Among them, Protenix showed the best performance with an accuracy of 80.8%, even surpassing AF3 which achieved 76.8%. When using MSAs (multiple sequence alignments), Chai-1 attained a success rate of 78.8%, compared to 70.7% without MSAs. Boltz exhibited similar success rate as Chai-1 without MSAs, at 71.7%. When the criteria were relaxed to medium (Fnat≥ 0.5), the success rates of the five methods increased to 85- 90%, with AF3 achieving the highest success rate of 89.9%. When the criteria were further lowered to acceptable (Fnat≥ 0.3), all methods utilizing MSAs reached success rates around 90%: AF3 at 90.9%, Chai-1 at 89.9%, and Protenix/Boltz-1 at 88.9%. Notably, while the use of MSAs in Chai-1 enhanced prediction accuracy, even without MSAs it still produced predictions comparable to other methods but has a significant efficiency advantage. (Figure 1A)

**Figure 1.**
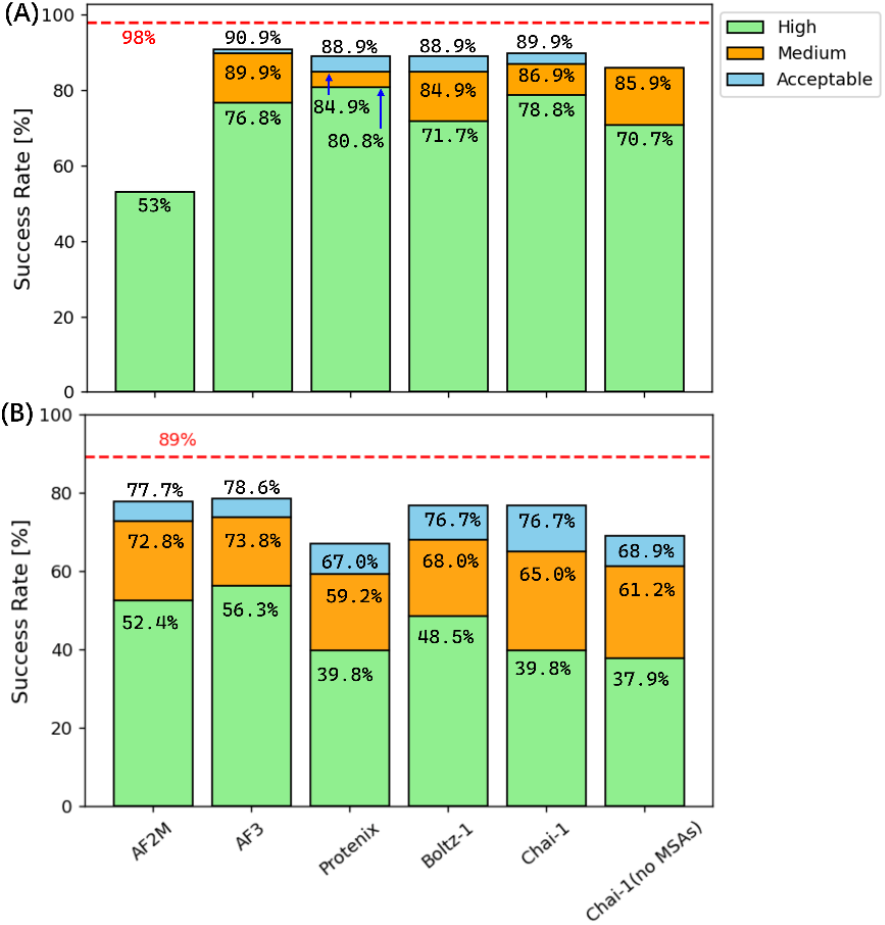
Comparison of success rates across different modeling methods. (A) the result of the first test set; (B) the results of the second test. Three criteria were used to define success: **High** (Fnat≥ 0.8), **Medium** (Fnat≥ 0.5), and **Acceptable** (Fnat≥ 0.3). The red line in the figure represents the theoretical success rate, i.e., selecting the best prediction from all models generated by all methods. Using this approach, the success rate under the acceptable criterion reaches 98% for the first test set, and 89% for the second test set.

As shown in Figure 1B, for the second test set, with high criteria, the success rates of all methods except AF2m decreased significantly. AF3 performed the best with 56.3%, while Protenix, Boltz-1, and Chai-1 had 39.8%, 48.5%, and 39.8%, respectively. When MSAs were not used, the success rate of Chai-1 decreased slightly to 37.9%. The prediction success rate of the new generation method for the second test set under the medium criteria and the acceptable criteria also decreased significantly, by 21% and 15% on average, respectively. The significant decrease in the success rate in the face of new structures indicates that the new generation modeling methods may generally have the risk of overfitting.

Representative results are illustrated in **Figure 2**, demonstrating the improved performance of next-generation modeling methods. **Figure 2A** displays the complex structure of the MUN domain of Munc13-1 with the membrane-proximal linker region (LR) of Syb2 (PDB ID: 6a30 [26]). The specific interactions between this protein receptor and the linker region short peptide coordinate SNARE complex assembly, facilitating neuronal exocytosis [26]. Munc13-1 is not a typical globular protein but adopts an elongated rod-like structure composed of multiple helices, lacking a conventional binding pocket. This makes binding site prediction challenging. While AF2m failed to correctly predict the short peptide’s binding site, other methods succeeded. **Figure 2B** shows the predicted complex structure of matrix metalloproteinase-2 (MMP-2) and its beta-amyloid precursor protein-derived inhibitor (a decapeptide). This inhibitor forms multiple long-range contacts or interactions with the protease to ensure binding specificity [27]. The challenge lies in the considerable diversity of the substrate-binding cleft within the MMP family, complicating the prediction of the correct binding mode. Unsurprisingly, all models for this system correctly predicted the binding site. However, AF2m failed to predict the positions of the first five peptide residues, while the positions of the latter five residues aligned well with the native structure. All next-generation methods accurately predicted the short peptide’s binding mode. **Figure 2C** depicts the prediction of rhomboid intramembrane proteolysis [28] and its substrate peptide. Similar to the previous example, AF2m identified the binding pocket but failed to predict the binding mode, placing the substrate further outward compared to its native position while all other methods accurately predicted the binding mode between the substrate and the protease.

**Figure 2.**
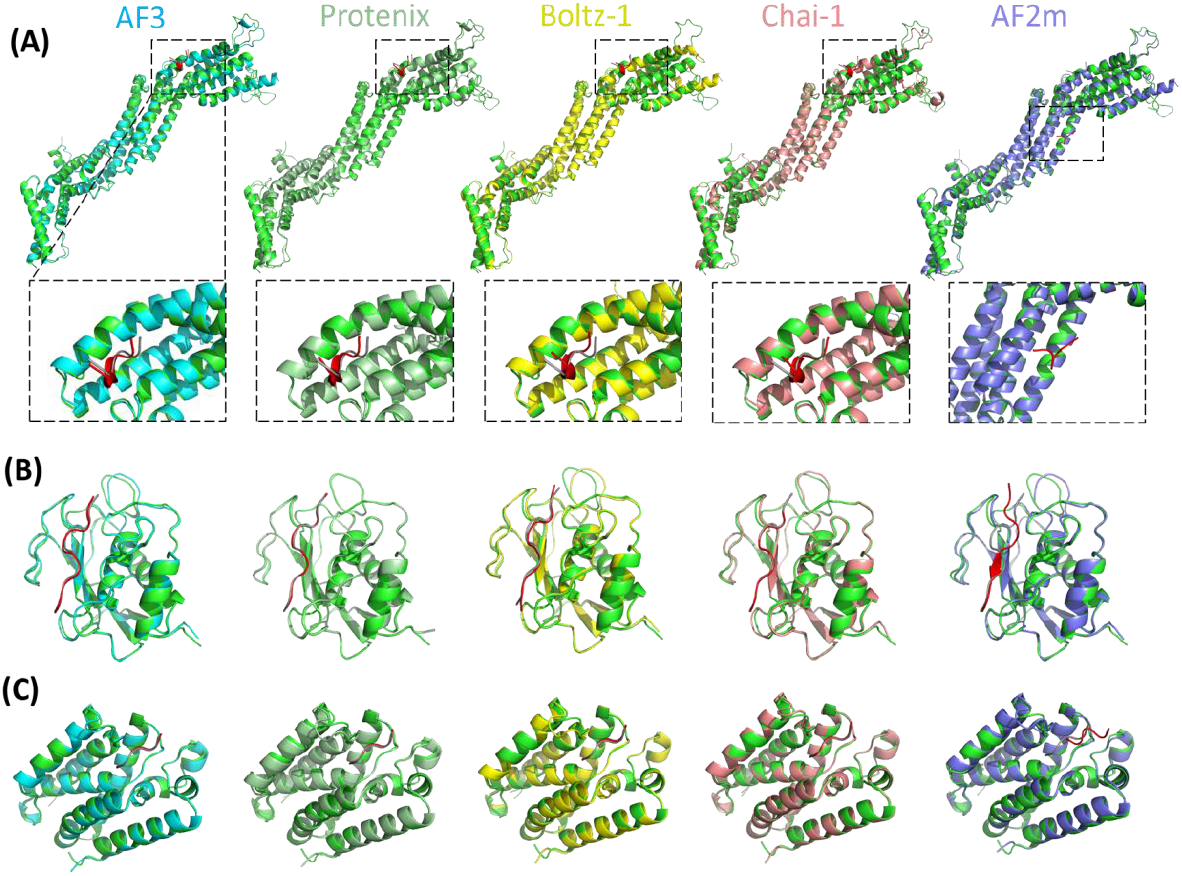
Representative cases demonstrating the superiority of next-generation modeling methods over previous-generation methods. **(A)** Modeling results for Munc13-1 system (PDBID: 6A30). **(B)** Modeling results for MMP-2 system (PDBID: 3AYU). **(C)** Modeling results for rhomboid intramembrane proteolysis system (PDBID: 6PJ8). In all panels, results from left to right correspond to AF3, Protenix, Boltz-1, Chai-1, and AF2m. The green structure represents the crystallographic conformation of the receptor, the gray structure denotes the crystallographic conformation of the ligand short peptide, and the red structure indicates the predicted conformation of the ligand short peptide.

However, there are still some challenges for all methods, even in the first test set. **Figure 3** summarizes three representative examples where these next-generation models underperformed. In **Figure 3A**, the crystal structure (PDBID: **3N2D**) [29] of a type I ribosome-inactivating protein complexed with a hexapeptide (SDDDMG) is compared with predictions. All four methods (AF3, Protenix, Boltz-1, and Chai-1) failed to predict the correct binding site, with inconsistent predicted binding positions across methods. Notably, Protenix also completely mispredicted the receptor protein structure. For the MATE multidrug transporter-MaL6 peptide complex (PDBID: **3WBN**) [30], all methods failed to identify the correct binding position (**Figure 3B**). A critical factor is that MaL6 is actually a cyclic peptide linked via a disulfide bond between the N-terminal and penultimate C-terminal CYS residues [31] yet we treated it as a linear peptide in all methods. Only Chai-1 predicted a beta-hairpin structure resembling the cyclic conformation (black box in **Figure 3B** right). Other models predicted extended peptide conformations with helical elements. We note that the MATE multidrug transporter contains multiple binding sites (shown in Figure 3B right, PDBID: 3VVR [32]), further complicating predictions. **Figure 3C** shows the crystal structure (PDBID: **5A29**) [33] and modeling results for Family 2 polysaccharide lyases (PL2s) complexed with their peptide partner. None of the four methods achieved Fnat≥ 0.8. Protenix produced the closest prediction to native (Fnat=0.71, DockQ=0.8). While AF3 identified the correct binding region, its peptide conformation deviated from native (Fnat=0.24, DockQ=0.37). Boltz-1 completely mispredicted the binding site. Although Chai-1 located the correct binding region, both the receptor conformation near the binding pocket and the peptide structure differed substantially from native.

**Figure 3.**
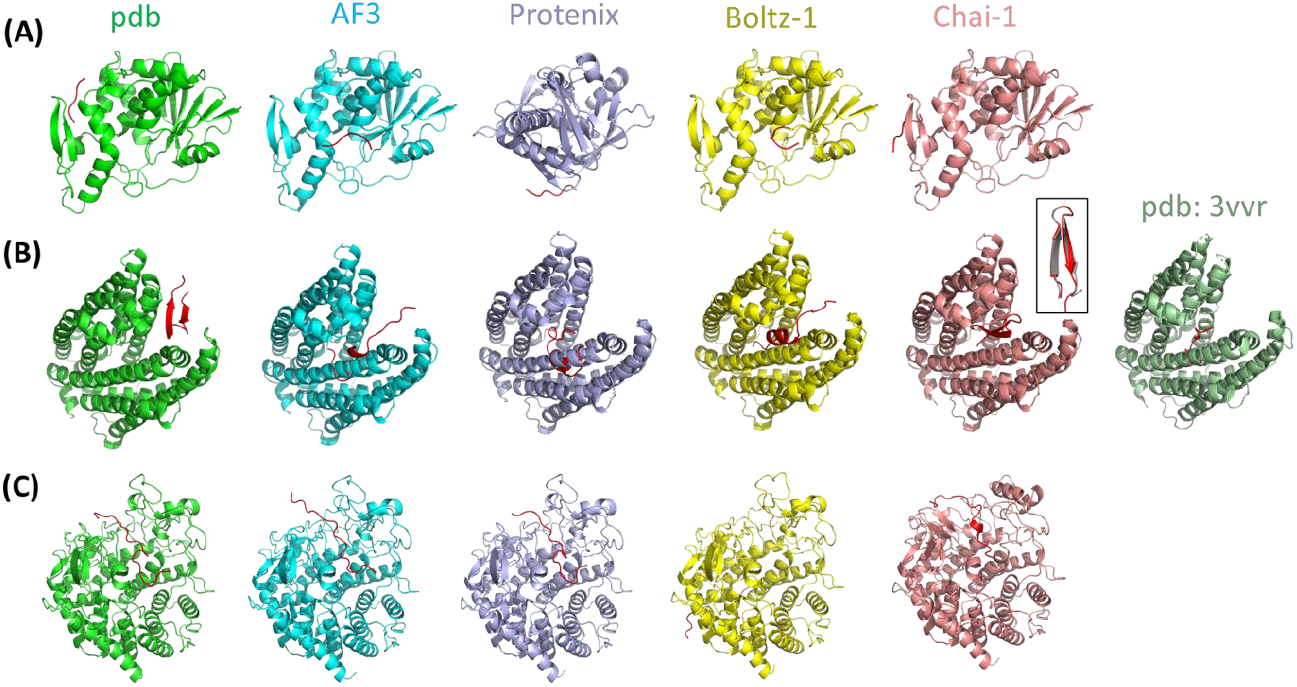
Representative cases of unsuccessful structural prediction. **(A)** Modeling results of type I ribosome-inactivating protein and hexapeptide (PDBID: **3N2D**). **(B)** Modeling results of MATE multidrug transporter-MaL6 peptide complex (PDBID: **3WBN**). **(C)** Modeling results of Family 2 polysaccharide lyases system (PDBID: **5A29**). In all subfigures, from left to right are the crystal structure, AF3, Protenix, Boltz-1, and Chai-1 results. The predicted structure of the red ligand peptide. The one at the far right of B is the crystal structure of another protein in the same family as **3WBN** (PDBID: **3VVR**), showing different binding sites.

Overall, the prediction failures can be categorized into three scenarios: 1. incorrect receptor protein structure prediction; 2. incorrect binding site prediction; 3. incorrect binding mode prediction. **Figure 4A, B** compares the distributions of TM-scores of receptors in the first and second test set across different methods. Overall, the AF3 method achieved the highest accuracy in receptor structure prediction, with an average TM-score of 0.97 for the first test set and 0.88 for the second test set. The average receptor TM-scores of 5 new generation methods are 0.94 for the first test set and 0.86 for the second test set. Compared with the significant decrease in the success rate of complex prediction, the performance of receptor structure prediction for the second test set, although it has also decreased, remains relatively consistent with that of the first test set. In particular, the number of predictions with low TM-scores (0-0.2) of the second test set is even lower than that of the first test set. The RMSD distribution also shows a similar trend: conformations below 2 angstroms are dominant in both test sets (**Figure S1 A**,**B**). This indicates that the challenge does not come from the receptor structure prediction. For comparison, the corresponding metrics of AF2m for the second test set were also calculated (**Figure S2**).

**Figure 4.**
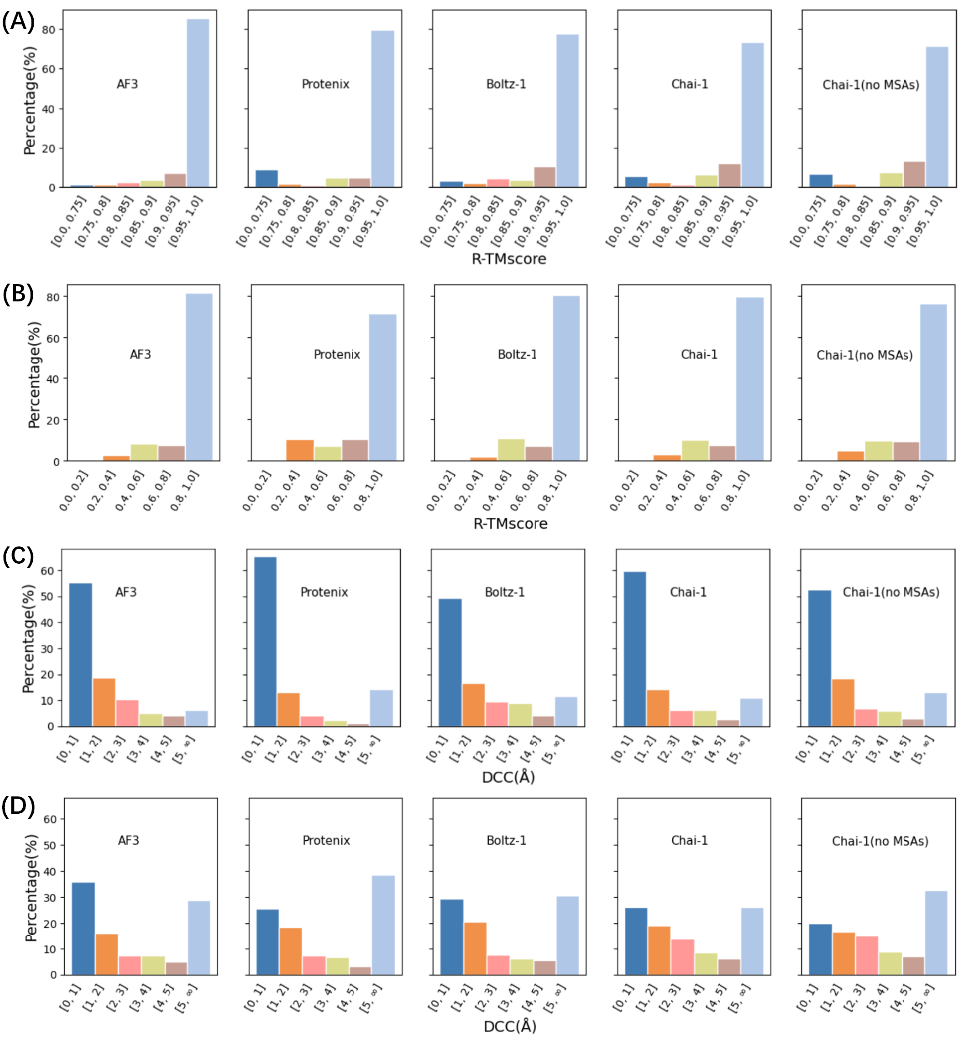
Comparison of prediction accuracy of receptors and pocket sites. **(A)** and **(B)** the TMscore distribution of receptor models relative to the native conformation for the first and second test set respectively. **(C)** and **(D)** the DCC distribution of predicted pocket sites for the first and second test set respectively.

We use DCA (the Distance from the Center of the predicted site to Any atom of the ligand) and DCC (the Distance from the Center of the predicted site to the Center of the ligand) to measure pocket prediction accuracy. DCA, widely used in prior studies [34-38], is the minimum distance between the predicted binding site center and any ligand atom. Peoples consider a predicted site correct if this distance is within 4 A. DCC, proposed by Skolnick and Brylinski (2008)[39], measures the minimum distance between the predicted binding site center and the ligand center, accounting for ligand size and providing higher success rates for larger ligands compared to DCA. DCC was recently used to compare Findsite and LIGSITEcsc [40]. As shown in **Figure 4C and D**, the performance of the first and second test sets differ significantly in terms of pocket prediction accuracy (measured by DCC). For the first test set, the percentage of DCC <2 A is 74.8%(AF3), 78.6%, (protenix), 66.3%(Boltz-1), 74.1%(Chai-1), 71.1%(Chai-1 no MSAs), while for the second test, the percentage of DCC <2 A is significantly reduced to 51.8%(AF3), 44.3%, (protenix), 50.1%(Boltz-1), 45.2%(Chai-1) and 36.9%(Chai-1 no MSAs), with an average decrease from 72.9% to 45.7%. In addition, the percentage of DCC > 5 A also increases significantly in the second test set, with an average increase from 11.1% to 31.1%. AF2m faces the same challenges as the new generation methods, as shown in the **Figure S2**. The DCA calculation results obtained the same conclusion as DCC (**Figrue S1 C, D**). When considering only pocket prediction performance, however, these structure prediction methods have become state-of-the-art (SOTA) in pocket prediction, surpassing deep-learning based algorithms like PointSite [35].

Structural modeling methods typically generate five models. These models vary in training parameters (number of templates/sequences used, training samples, and training time), producing five structural models for each input sequence. As shown in **Figure 5** (and **Figure S3**), we examine the performance of these five models. In all methods, model ranked_0 has the highest probability of being the best model (**Figure 5A** and **Figure S3A**). Notably, Protenix’s ranked_0 model has a 57% chance of being the best prediction. However, the differences among the five models are not significant, as their prediction success rates are similar, and ranked_0 is not always the most successful model (**Figure 5B** and **Figure S3B**).

**Figure 5.**
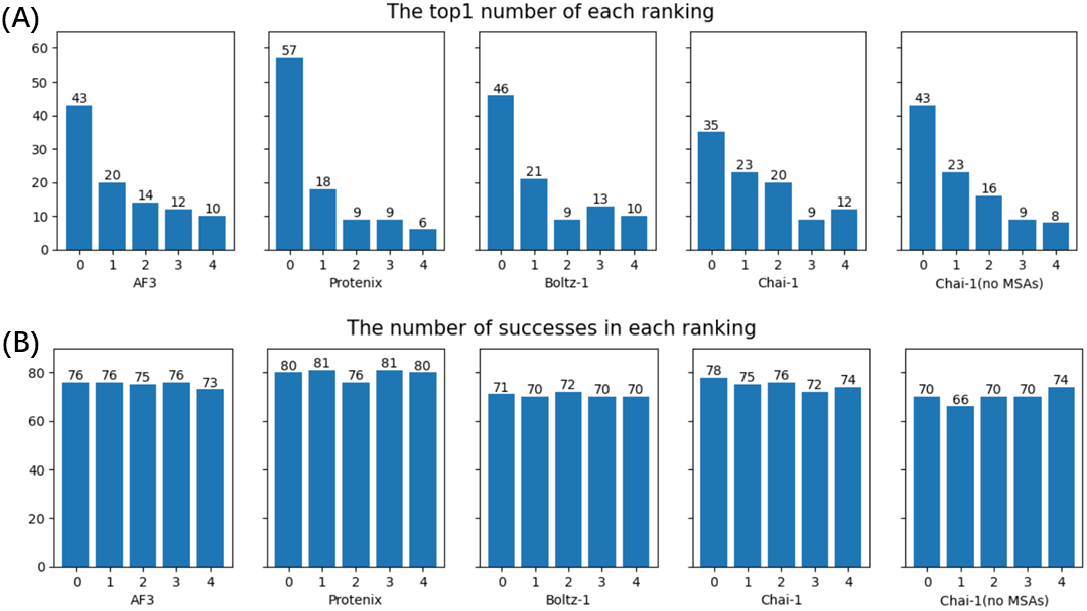
Performance of each model (from ranked 0 to ranked 4) of each method for the first test set. **(A)** The probability of each model being the best prediction among the other four. **(B)** Success rate (Fnat≥ 0.8) of each model.

To pick the most likely successful prediction from 5 models (and between different methods), we checked the relationships between multiple indicators related to the predicted model (such as pLDDT, ipTM, RMSD in molecular dynamics simulation, and the number of interactions between peptides and protein receptors) and the deviation from the native structure (by Fnat, and DockQ). DockQ is a metric for assessing the quality of protein - protein docking models [23]. It combines Fnat, LRMSD (ligand RMSD) and iRMSD (interface residue RMSD) to give a continuous score in [0.0, 1.0], where a higher value means better accuracy. DockQ’s advantage lies in its continuous scoring (rather than putting models into different quality categories), which helps analyze model similarity and prediction performance in more detail.

For the first test set, when pLDDT is under 70, Fnat stays below 0.6 and DockQ below 0.5. Most points cluster in regions where pLDDT ≥ 90 with Fnat≥ 0.8 or DockQ≥ 0.6 (left panels of **Figure 6A,B**). High-quality predictions are more scattered in ipTM scores than pLDDT, mostly above ipTM ≥ 0.5. MD-derived RMSD correlates with Fnat and DockQ (middle panels of **Figure 6A,B**). RMSD, calculated using the predicted structure as reference, indicates greater deviation and instability when larger. Few high-quality predictions occur when RMSD exceeds 6 A, while most are high-quality when RMSD is below 2 **Figure S4A,B**(middle). Though more interactions intuitively suggest a firmer binding, total interactions in **Figure S4A,B**(left) don’t strongly correlate with Fnat and DockQ. High-quality predictions are scattered in areas with fewer interactions, and points with the most interactions don’t correspond to high Fnat and DockQ scores. Note: To account for peptide length’s impact on interaction numbers, #PPI in the figure is PPI count divided by peptide length, i.e., average interactions per residue. Unlike the first test set, for the second test set, the correlations of all the above metrics with those of DockQ or Fnat become worse, and the distribution of points is more scattered (**Figure 6C,D** and **S5**).

**Figure 6.**
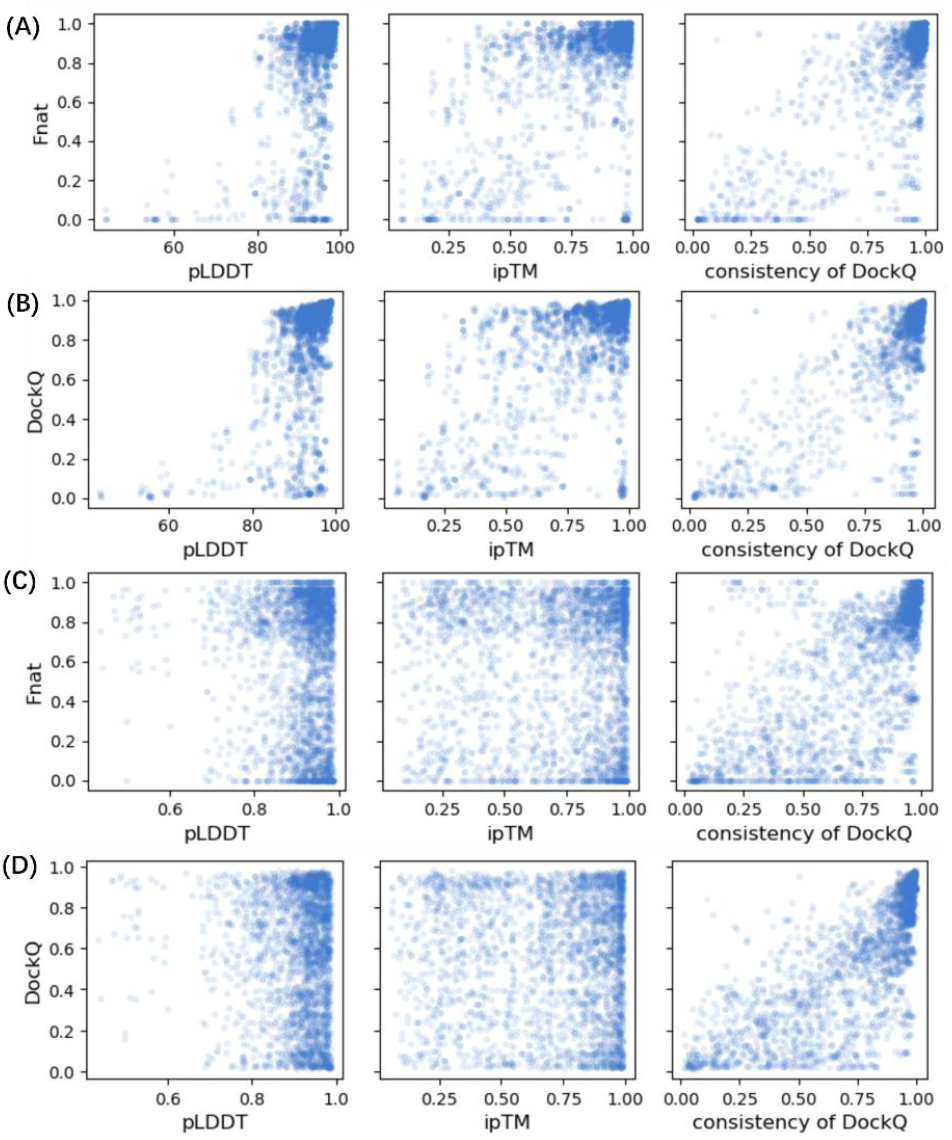
Correlation of different indicators with Fnat and DockQ. These indicators include: pLDDT of the entire complex (left), ipTM of the peptide (middle), and consistency using DockQ as indicator (right). **(A)** and **(B)** for the first test set. **(C)** and **(D)** for the second test set.

In addition to the above indicators, we introduced the “consistency” to select the best predictions from five methods and each method’s five models. We calculate DockQ for the five models within the same method, using each other as references. The average DockQ of a model with the other four (using the models as references) is its consistency score. **Figure 6** (right) shows that the consistency of DockQ (using other models as references) is most significantly correlated with Fnat and DockQ (using native as reference) for the first and second test set. That means when a method gives high-quality predictions, its five models are usually consistent. We also check the consistency by counting identical interactions (same participating atom pairs and interaction types). Similar to #PPI, this indicator generally shows no correlation with Fnat and DockQ (**Figure S4** and **S5**). However, it is useful for model screening (see discussion below), probably because it essentially reflects the consistency of the model rather than the number of interactions.

We tried to pick the best predictions from all methods and models using the above indicators. **Figure 7, S6** and **S7** summarize the success rate of the selected models. For the first test set, most indicator-selected predictions perform better (except for #PPI) than any single-method predictions. The high-quality prediction success rate increased from 81% (Protenix) to a maximum of 89% (selected by DockQ consistency). For the second test set, the consistency indicator can help, and the models selected by DockQ consistency can obtain a success rate of 60.2%, which is slightly better than AF3’s 56.3%. The other indicators did not play a positive role.

**Figure 7.**
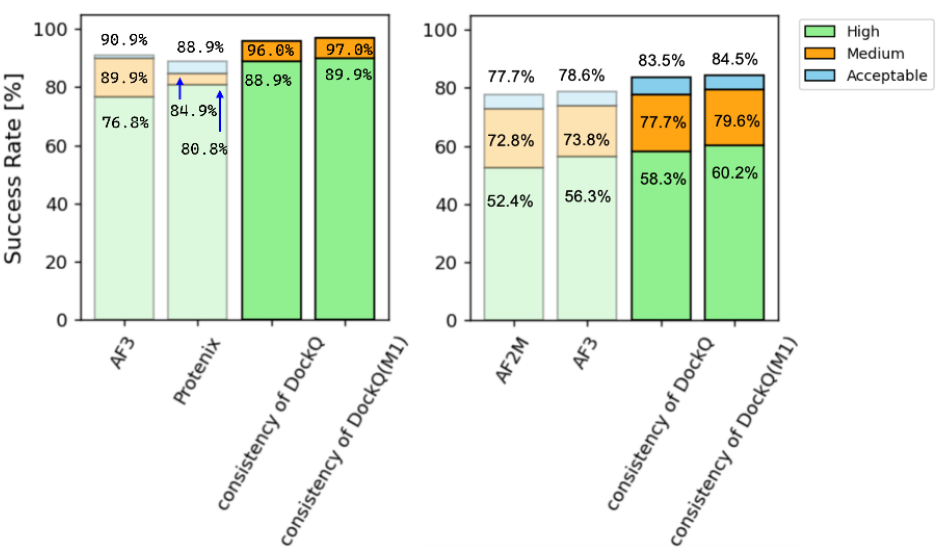
Success rates of protein-peptide structural predictions after filtering methods/models with indicators. Left and right panels are the results of the first and second test sets respectively. Standards for High, Medium, and Acceptable are the same as in Figure 1. The label “M1” represent AF3 + Protenix combination for the first test set, and AF2m + AF3 combination for the second test set respectively. The combinations are used as model pools and are screened by consistency of DockQ.

Running multiple modeling methods for the same system and selecting the best predictions with consistency indicators can improve the success rate of protein-peptide complex structure prediction, indicating some complementarity among methods. As shown in **Figure 8A** (and **S8**), for the first test set, 52 systems were successfully predicted by all methods. It is worth noting that 7 models were predicted successfully only by Protenix, and 2 models were predicted successfully only by Chai-1. When combining two methods, higher success rates can be achieved. AF3 and Protenix had 91 successful predictions together, as did Protenix and Chai-1. The combination of AF3, Protenix, and Chai-1 had 94 successful predictions. For the second test set (**Figure 8B** and **S9**), only 24 systems were successfully predicted by all new generation methods. The overlap between methods was significantly reduced compared to the first test set.

**Figure 8.**
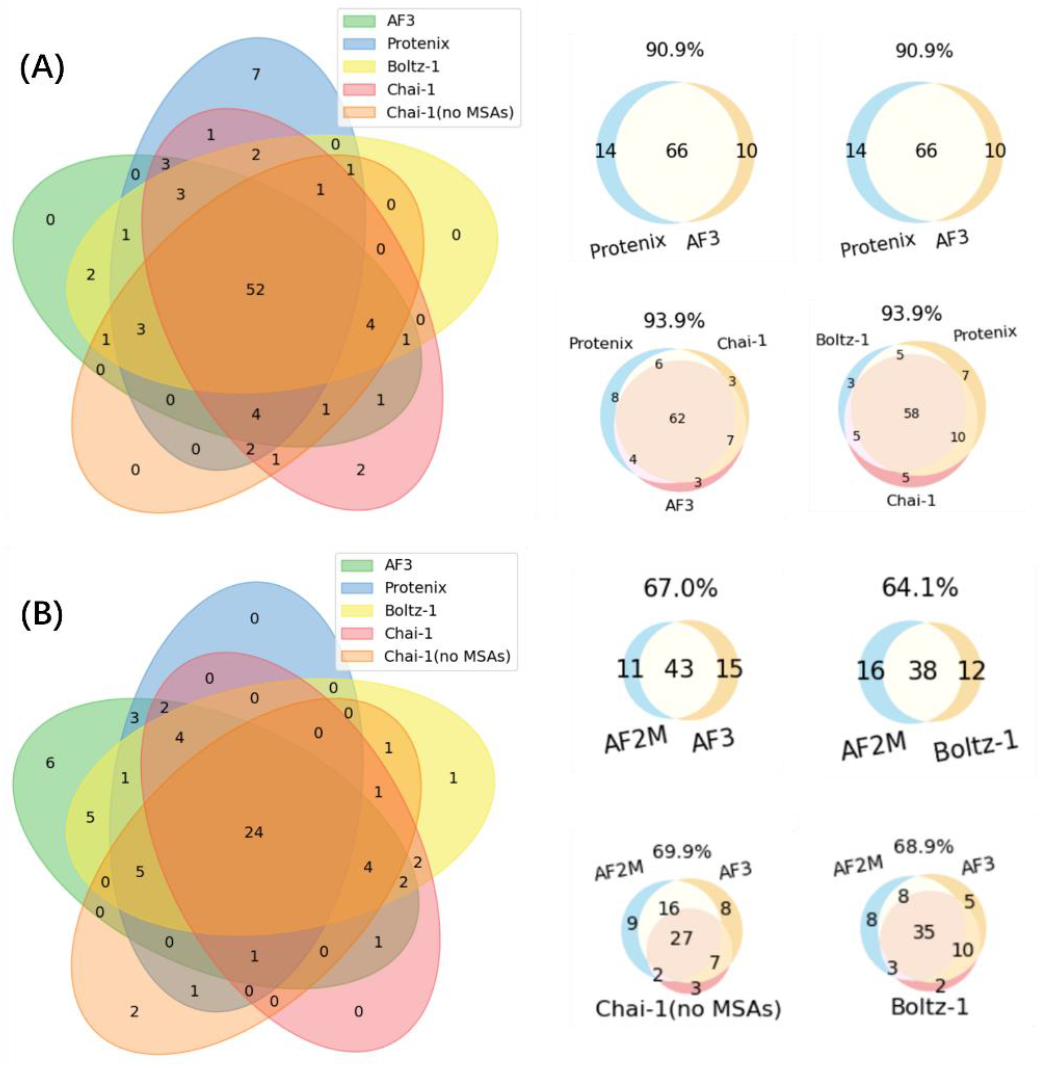
Overlap of successful cases predicted by different modeling methods. **(A)** Overlap of methods with each other in the first test set. **(B)** Overlap of methods with each other in the second test set. Each panel shows the overlap of five models on the left and shows the best combinations on the right.

Combining all methods as model pools and selecting models based on DockQ consistency, the success rates improved as mentioned earlier. However, we can also achieve a high success rate by selecting only some of the methods and combining them. As shown in the **Figure 7**, screening the AF3 and Protenix combination with consistency yielded an 89.9% high-quality prediction success rate for the first test set. screening the AF3 and Af2m combination with consistency yielded a 59.2% high-quality prediction success rate for the second test set. Compared with using the combination of all methods, using the combination of two methods can achieve the best prediction effect. This may be partly due to the fact that too many methods increase the difficulty of selecting the best confirmation. This conclusion has high guiding value in practical applications.

## Method

### Data Collection and Processing

The second test set was retrieved from the RCSB Protein Data Bank (PDB) using the following selection criteria: 1. Structures released after September 30, 2021, to ensure temporal separation from the model’s training set; 2. Entries containing peptides with chain lengths between 3-50 amino acids paired with proteins shorter than 900 residues; 3. Systems with less than 40% sequence similarity as determined by CD-HIT clustering (v4.8.1) [41] using standard parameters. A final quality control step was implemented to select structures demonstrating high consistency between experimental electron density maps and corresponding amino acid sequences (resolution ≤ 3.0 A, R-factor ≤ 0.25). This curation process aimed to minimize potential discrepancies between input sequences and structural data that might confound model predictions. The final test set contained 91 protein-peptide complexes.

### Structure Prediction

We performed protein-peptide structure prediction using AF3 (version 3.0.0) with all default parameters, generating 5 predicted models through 1 random seed. For multiple sequence alignments (MSAs), the Jackhmmer module from HMMER [42] (version 3.3.2) was employed. The Protenix version used in this work was 0.3.7, with runtime parameters consistent with those specified in the “inference_demo.sh” script from its source code repository (https://github.com/bytedance/Protenix). Each prediction produced 5 models, with diffusion model steps set to 200, 10 cycles of backbone inference iterations, and a random seed fixed at 101. Following the AF3 protocol, we utilized the Jackhmmer module in HMMER (version 3.3.2) to construct MSAs for Protenix. For Boltz-1 (version 0.4.0), structure predictions were conducted using default parameters, generating 5 models per run. MSA construction was automatically completed through the mmseqs2 service [43]. The Chai-1 framework (version 0.5.1) was implemented with 5 trunk recycling iterations and 200 diffusion model steps, producing 5 models per prediction. The Chai-1 model offers optional MSA usage, where MSAs are automatically generated through the mmseqs2 service. When MSAs were not employed, the large language model ESM [44] (Evolutionary Scale Modeling) was substituted as an alternative approach.

### Interaction Analysis

To analyze interaction methods, we employed PLIP [45](version 2.4.0), a tool designed for characterizing non-covalent protein-ligand (short peptide) interactions in 3D structures. Our analysis primarily focused on four interaction types: hydrogen bonds, salt bridges, pi-stacking, and pi-cation interactions. The criteria were defined as follows. For hydrogen bonds, All pairs of hydrogen bond acceptors and donors were identified with donor hydrogen atoms and acceptor atoms within 0.5–4.1 A distances and donor angles exceeding 100 degrees. For salt bridges, protein-peptide residue pairs with opposite charges were considered if their centroid distance was less than 5.5 A. pi-Stacking interactions were counted when aromatic rings were arranged in parallel or perpendicular orientations, with centroid distances below 5.5 A and angular deviations of ring planes under 30 degrees. For pi-Cation, all aromatic ring-cation pairs were included if their distance was within 6 A.

### Molecular dynamics

The initial structures used in the simulations are the predicted models. For each system, the protonation state of neutral HIS residues were automatically determined by the pdb2gmx tool from GROMACS [46]. The DES-Amber [47] force field was applied to proteins, and the TIP4P-D[48] model was utilized for water molecules. The simulation box was set to be a cube with a boundary of 10 A, and a 0.1 M NaCl buffer solution was added to ensure electrical neutrality of the system. All simulations were conducted using OpenMM[49] (version 7.7). Each system was minimized for 1000 steps followed by a 500 ps pre-equilibration period. The simulations were run using the Monte Carlo barostat method at 1 atm for 10 ns, with MD snapshots saved every 25 ps. The integrator chosen was the Langevin integrator with a temperature of 320 K and an integration timestep of 4 fs. The masses of hydrogen atoms and adjacent heavy atoms on the protein were redistributed so that the total mass of the chemical group remained unchanged, while the mass of hydrogen atoms was increased to 4 atomic mass units to reduce the frequency of angle bending motions associated with hydrogen, thereby increasing the integration time step.

## Conclusion

This study systematically evaluated the performance of next-generation structure prediction methods (including AlphaFold3, Protenix, Chai-1, and Boltz-1) in modeling protein-short peptide complexes on two benchmark test sets. The results for the first test set demonstrate that compared to previous methods (such as AlphaFold2-multimer), these new approaches significantly improved the prediction success rate under stringent criteria (Fnat≥ 0.8) to 70-80%, with Protenix achieving the highest accuracy of 80.8%. However, for the second test set (which excluded structures included in the training set of all methods), the success rate of the next-generation modeling methods (with Fnat≥ 0.8) dropped significantly to 40-65%, suggesting that the next-generation modeling methods are at risk of overfitting. For both test sets, some failure cases persist. These failures primarily stem from errors in protein receptor structure prediction, binding site misidentification, or incorrect binding mode prediction. Correlation analysis demonstrated that an important reason for the reduced success rate was the failure to predict the binding sites of the”new”complexes. We further investigated the relationship between various indicators—such as pLDDT, ipTM, RMSD, the number of interactions between peptides and protein receptors (#PPI), model consistency—and prediction accuracy (measured by Fnat and DockQ). Notably, we found that the consistency among five models predicted by a single method serves as a robust metric for selecting the optimal model. Overlap analysis of successful predictions from different methods revealed complementary advantages between approaches. By combining two methods or three methods and employing consistency metrics for selection, the success rate could be further increased for both test sets. Overall, this study not only validates the superiority of next-generation structure prediction methods in protein-short peptide complex modeling but also provides critical references and directions for future research.

