## supplemental information for "Benchmarking AlphaFold3-like Methods for Protein-Peptide Complex Prediction"

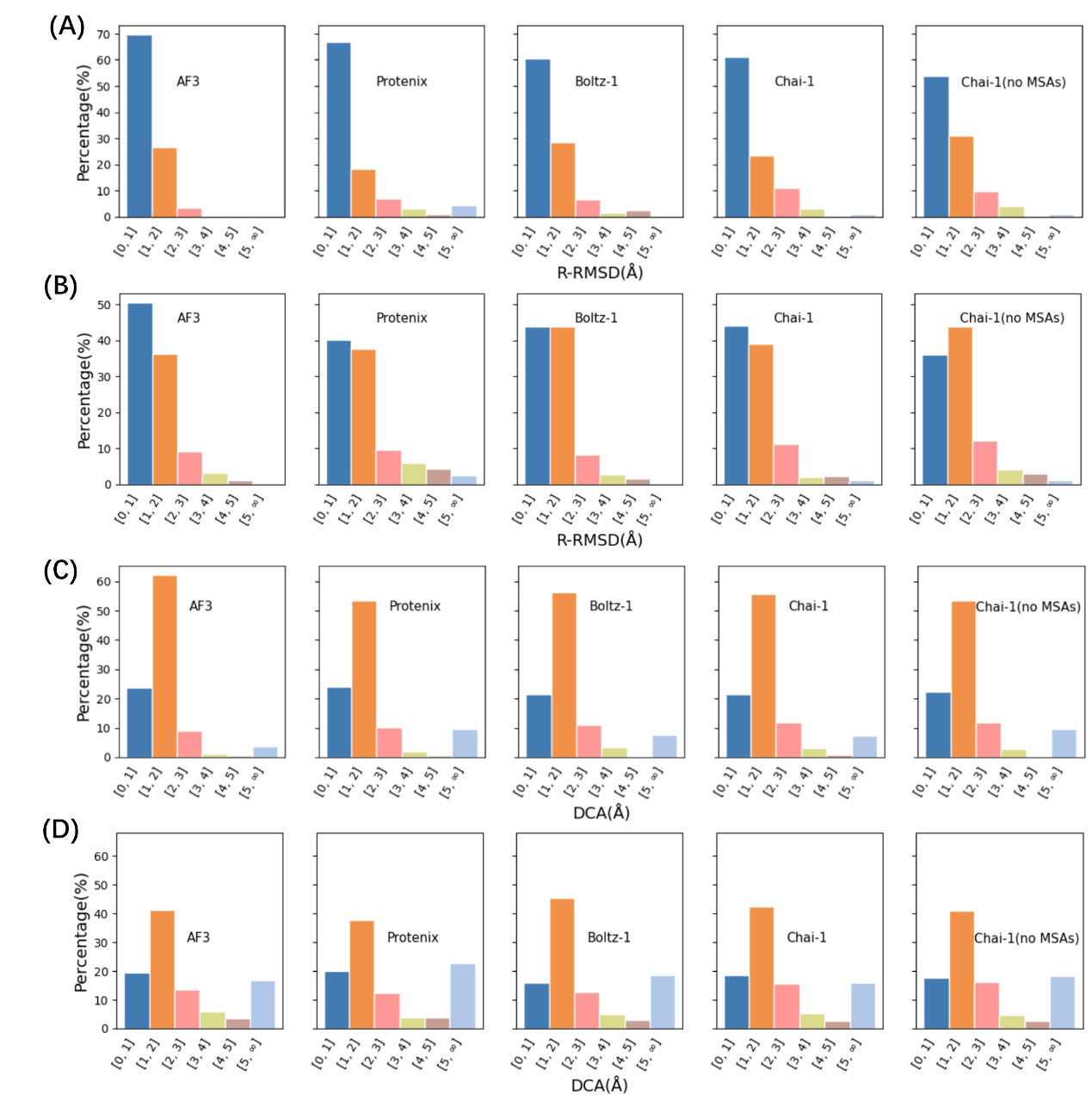

**Figure S1**. Comparison of prediction accuracy of receptors and pocket sites. **(A)** and **(B)** RMSD distribution of receptor models predicted by five methods relative to the native conformation (All CA atoms of residues were selected for RMSD calculation) for the first and second test set respectively. **(C)** and **(D)** DCA distribution of predicted pocket sites for the first and second test set respectively.

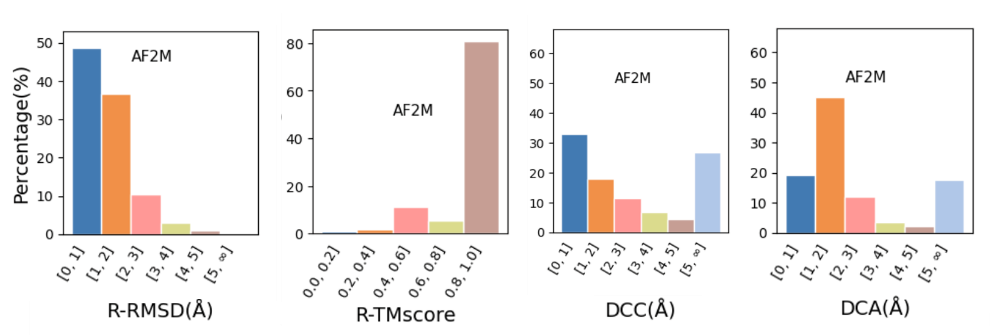

**Figure S2**. The prediction accuracy of receptors and pocket sites of AF2m

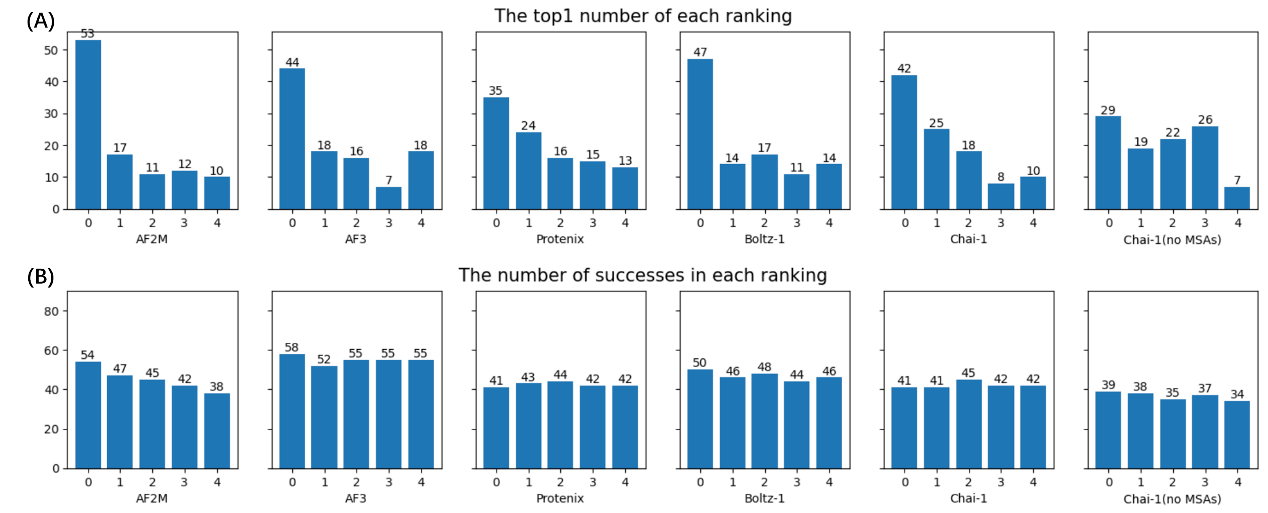

**Figure S3.** Performance of each model (from ranked 0 to ranked 4) of each method for the second test set. **(A)** The probability of each model being the best prediction among the other four. **(B)** Success rate (Fnat≥ 0.8) of each model.

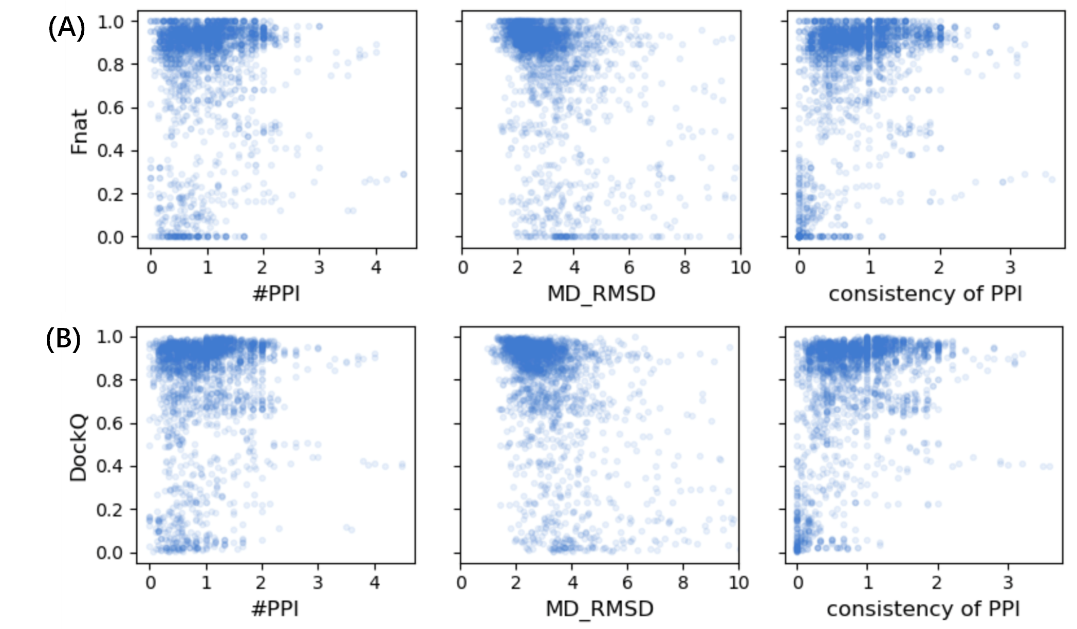

**Figure S4.** Correlation of different indicators with Fnat and DockQ in the first test set. These indicators include: average number of interactions (#PPI) between proteins and short peptides per residue, RMSD changes from MD simulations (using the predicted structure as the reference), and consistency using identical #PPI count as indicator.

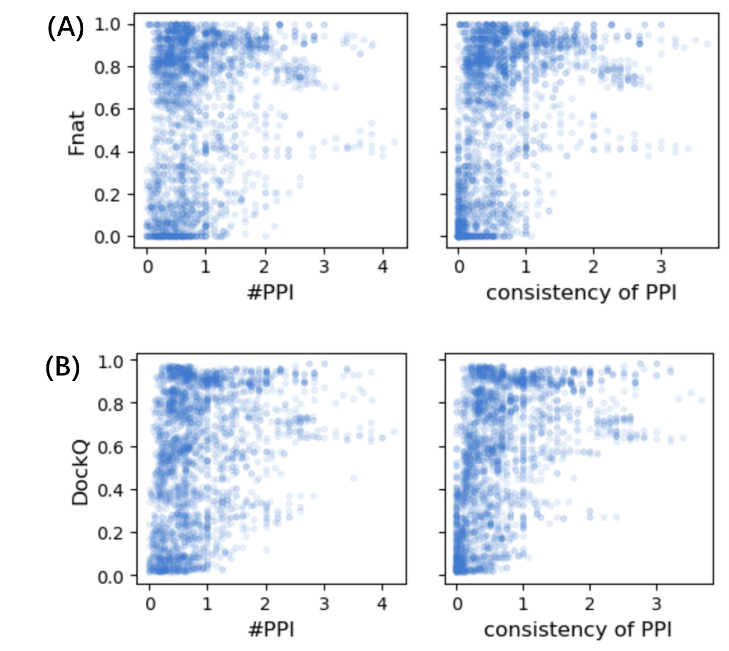

**Figure S5.** Correlation of different indicators with Fnat and DockQ in the second test set. These indicators include: average number of interactions (#PPI) between proteins and short peptides per residue, and consistency using identical #PPI count as indicator.

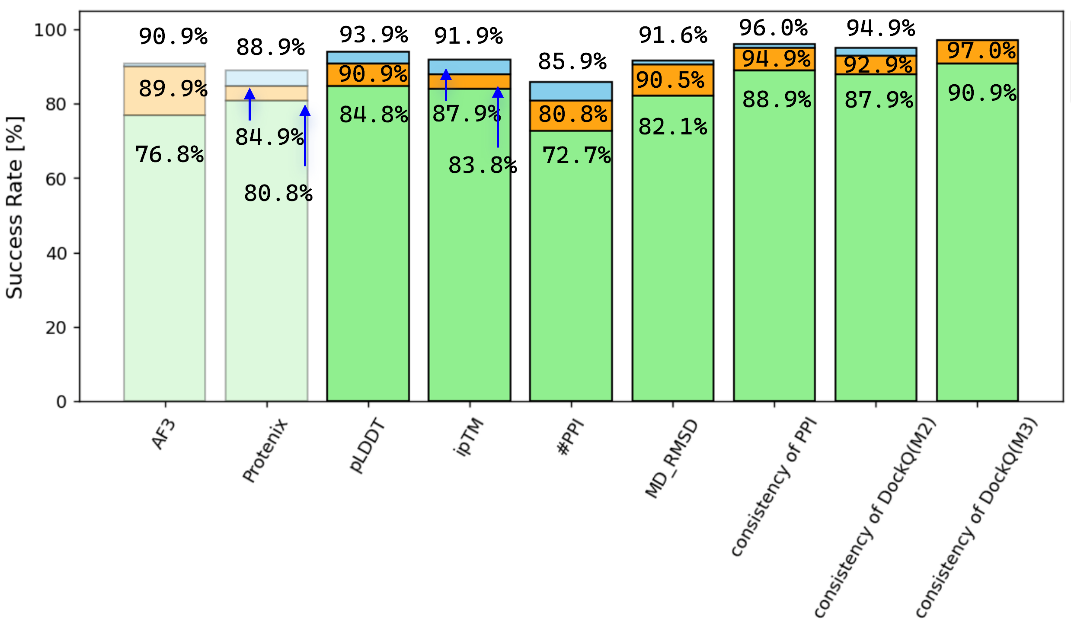

**Figure S6.** Success rates of protein-peptide structural predictions after filtering methods/models with indicators for the first test set. Standards for High, Medium, and Acceptable are the same as in Figure 1. The label “M2” represent AF3 + Protenix combination for the first test set, and AF2m + AF3 combination for the second test set respectively. The label M2, M3 in the two bars on the right represent Chai-1 + Protenix, and AF3 + Chai-1 + Protenix combination respectively. The combinations are used as model pools and are screened by consistency of DockQ.

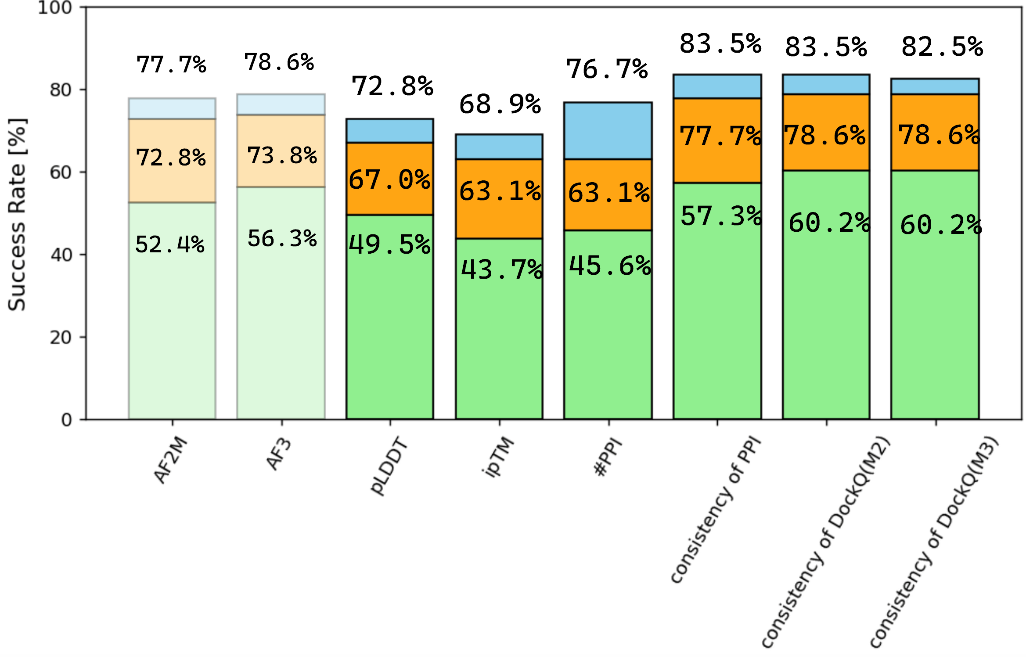

**Figure S7.** Success rates of protein-peptide structural predictions after filtering methods/models with indicators for the second test set. Standards for High, Medium, and Acceptable are the same as in Figure 1. The label M2, M3 in the two bars on the right represent AF2m + AF3 + Chai-1 (no MSAs), and AF2m + AF3 + Boltz-1 combination respectively. The combinations are used as model pools and are screened by consistency of DockQ.

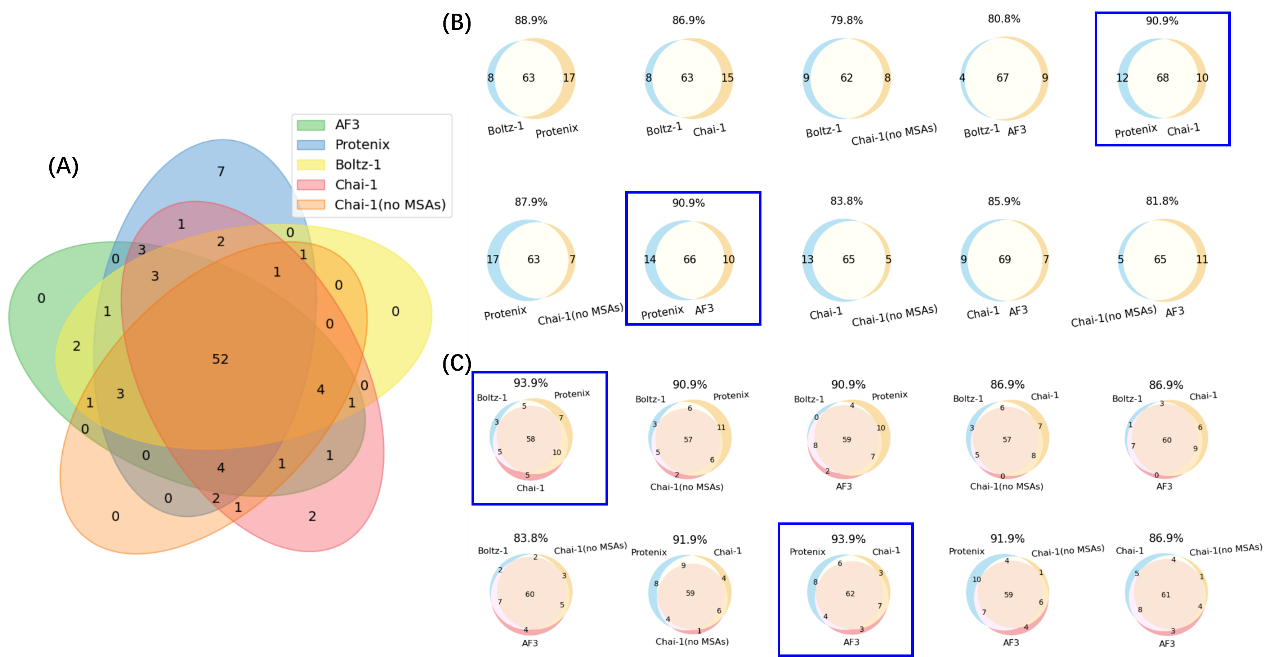

**Figure S8.** Overlap of successful cases predicted by different modeling methods for the first test set. **(A)** Overlap of the five methods with each other. **(B)** Overlap of successful cases in two-method combination.. **(C)** Overlap of successful cases in three-method combination.

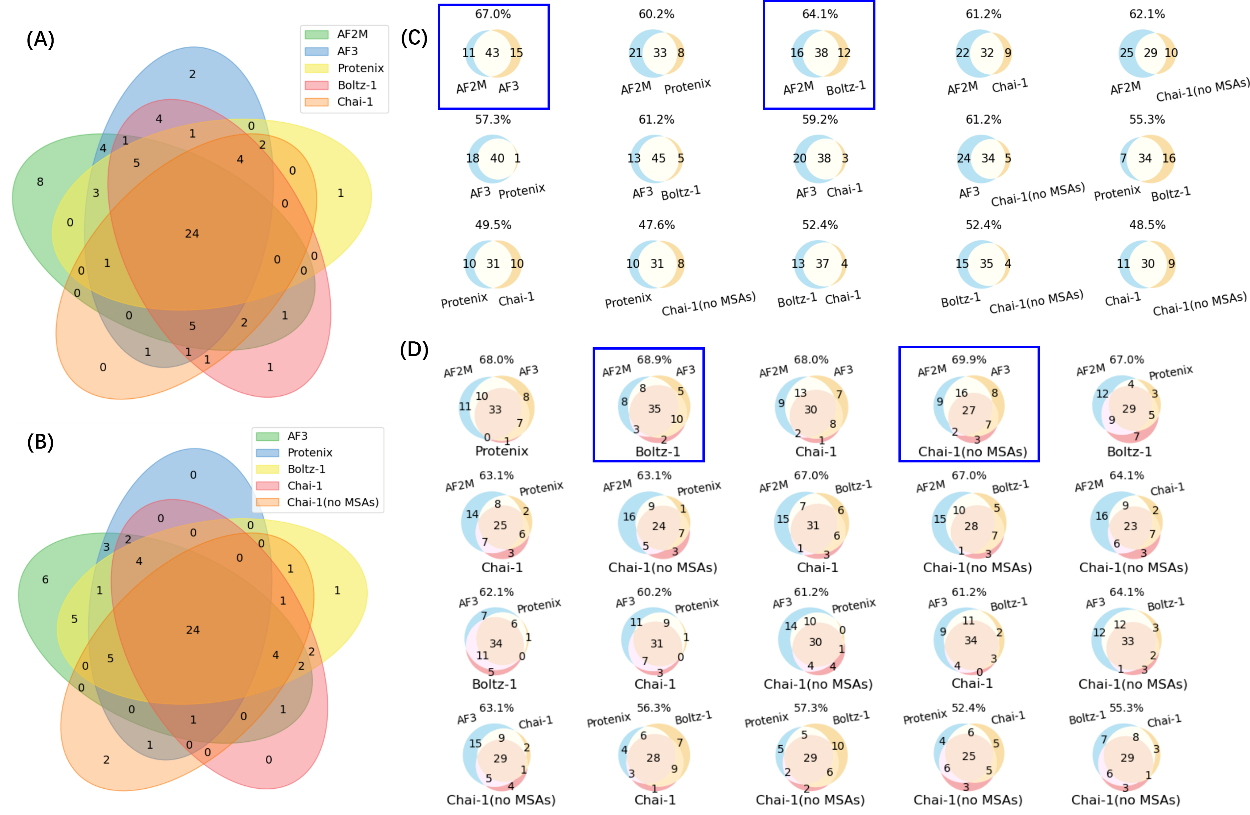

**Figure S9.** Overlap of successful cases predicted by different modeling methods for the second test set. **(A)** Overlap of the five methods (AF2m, AF3, Protenix, Boltz-1, Chai-1) with each other. **(B)** Overlap of the five methods (AF3, Protenix, Boltz-1, Chai-1, Chai-1 no MSAs) with each other. **(C)** Overlap of successful cases in two-method combination. **(D)** Overlap of successful cases in three-method combination.
